# ATXN2L primarily interacts with NUFIP2, the absence of ATXN2L results in NUFIP2 depletion, and the ATXN2-polyQ expansion triggers NUFIP2 accumulation

**DOI:** 10.1101/2025.03.06.641895

**Authors:** Jana Key, Luis-Enrique Almaguer-Mederos, Arvind Reddy Kandi, Nesli-Ece Sen, Suzana Gispert, Gabriele Köpf, David Meierhofer, Georg Auburger

**Author notes:** joint first authorship (Jana Key, Luis-Enrique Almaguer-Mederos). Correspondence to Dr. Georg Auburger.

## Abstract

The cytoplasmic Ataxin-2 (ATXN2) protein associates with TDP-43 in stress granules (SG) where RNA quality control occurs. Mutations in this pathway underlie Spinocerebellar Ataxia type 2 (SCA2) and Amyotrophic Lateral Sclerosis. In contrast, Ataxin-2-like (ATXN2L) is predominantly nuclear, more abundant, and essential for embryonic life. Its sequestration into ATXN2 aggregates may contribute to disease. In this study, we utilized two approaches to clarify the roles of ATXN2L. First, we identified interactors through co-immunoprecipitation in both wild-type and ATXN2L-null murine embryonic fibroblasts. Second, we assessed the proteome profile effects using mass spectrometry in these cells. Additionally, we examined the accumulation of ATXN2L interactors in the SCA2 mouse model, Atxn2-CAG100-KnockIn (KIN). We observed that RNA-binding proteins, including PABPN1, NUFIP2, MCRIP2, RBMS1, LARP1, PTBP1, FMR1, RPS20, FUBP3, MBNL2, ZMAT3, SFPQ, CSDE1, HNRNPK, and HNRNPDL, exhibit a stronger association with ATXN2L compared to established interactors like ATXN2, PABPC1, LSM12, and G3BP2. Additionally, ATXN2L interacted with components of the actin complex, such as SYNE2, LMOD1, ACTA2, FYB, and GOLGA3. We noted that oxidative stress increased HNRNPK but decreased SYNE2 association, which likely reflects the relocalization of SG. Proteome profiling revealed that NUFIP2 and SYNE2 are depleted in ATXN2L-null fibroblasts. Furthermore, NUFIP2 homodimers and SYNE1 accumulate during the ATXN2 aggregation process in KIN 14-month-old spinal cord tissues. The functions of ATXN2L and its interactors are therefore critical in RNA granule trafficking and surveillance, particularly for the maintenance of differentiated neurons.

## Introduction

The quality control of ribonucleoprotein (RNP) granules during long-distance transport in neurites is crucial for the stimulus-dependent mRNA translation in synapses [1–5]. Impairments in this pathway cause neurodegeneration preferentially of motor neurons and cerebellar neurons [6–14]. When cellular growth periods are interrupted by significant damage, the RNP granules coalesce into cytosolic stress granules (SG) where surveillance mechanisms decide to repair individual RNAs or degrade them in P-bodies [15–17]. The trafficking and surveillance machinery controls association with or dissociation from mRNAs at adenine-enriched sequences, known as poly(A) stretches [18], or at uracil-enriched sequences, known as AU-rich elements (ARE) [19]. It also controls the RNA winding into double-strand (ds)-hairpins or their unwinding via helicases [20], assesses the RNA posttranscriptional modifications [21], and liquid-liquid phase separation [22].

As a prime member of this SG surveillance machinery [23–26], Ataxin-2 contains a Like-Sm (LSm) domain together with an LSm associated domain (LSm-AD) sequence to interact with helicases such as DDX6 [23, 25, 27–29] and to bind single-stranded (ss) ARE [30, 31]. Furthermore, it contains a PAM2 motif to associate with ss-poly(A) stretches indirectly via poly(A)-binding-proteins [32–37]. These motifs enable not only surveillance of mRNAs, but also effects on rRNA, microRNA and long-noncoding RNA effects have been documented [38–42]. This characteristic combination of sequence motifs is first detected in eukaryotic organisms, such as in the yeast ortholog PBP1 [43–45], and exists as a single copy in nematodes such as *Caenorhabditis elegans* ATX-2 [46–50], or flies such as *Drosophila melanogaster* dATX2 [51–53]. In contrast, land plants contain two gene copies named *CID3* versus *CID4* [37, 54, 55], while mammals also contain two gene copies known as the less abundant *ATXN2* versus the more abundant *ATXN2L*, with complementary roles for translation oscillations and SG [56, 57]. Under unstressed growth conditions, Ataxin-2 isoforms are localized with the translation apparatus at the endoplasmic reticulum [34, 58–63] and with microtubules [64–67].

ATXN2 was first described, because pathogenic expansions of an N-terminal polyglutamine (polyQ) stretch from the normal length Q22/Q23 beyond a size of Q33 cause the autosomal dominantly inherited, neurodegenerative disorder Spinocerebellar Ataxia type 2 (SCA2) [68–71], where the glutamatergic corticospinal motor neurons and the glutamatergic cerebellar granular neuron projections to Purkinje cells are preferentially affected [72–81]. Much more infrequently, a Parkinson’s disease phenotype is triggered by the same monogenic polyQ expansions due to early loss of dopaminergic midbrain neurons [82–88]. ATXN2 polyQ expansions of intermediate size between Q27 and Q33 act as polygenic modifiers that exacerbate the progression of the adult motor neuron diseases Amyotrophic Lateral Sclerosis (ALS) and Fronto-Temporal Lobar Dementia (FTLD) [9, 89–96], and of Parkinson syndrome (Parkinson plus) [97, 98]. Conversely, the knockout (KO) or knockdown (KD) of ATXN2 was shown to strongly mitigate the progression of such diseases [12, 99, 100], and this makes ATXN2 a prime pharmacological target for the prevention of neurodegenerative processes that are triggered by RNA toxicity [101–104].

Although ATXN2L is the more abundant and more important paralog in cellular life, and although it may play a compensatory role for ATXN2 mutations [105], only a few studies have focused on it so far. Mice with homozygous deletions of *Atxn2l* exons 5-8, which results in the production of an N-terminal fragment but absence of the LSm, LSm-AD and PAM2 domains until the C-terminus, show mid-embryonic lethality with brain cortex lamination defects, while the heterozygous mutants appeared normal [106]. In comparison, the less abundant ATXN2 paralog would appear barely relevant, since mice with homozygous deletions of *Atxn2* exon1, which results in the production of a minimal N-terminal fragment but absence of the LSm, LSm-AD and PAM2 domains until the C-terminus, show only age-associated obesity, hepatosteatosis, insulin resistance and dyslipidemia, with reduced fertility and locomotor hyperactivity [58, 60, 61, 107, 108]. ATXN2L and ATXN2 expression is conversely modulated by growth factor signaling versus stress conditions [61, 106, 109]. Unlike ATXN2, the human promoter for ATXN2L expression is shown to change conformation upon exposition to G-quadruplex ligands [110]. Again unlike ATXN2, the human transcript for ATXN2L contains a SINE-VNTR-Alu element, which could modulate exon extension or alternative transcript regulation by stress events [111]. An intron in the ATXN2L pre-mRNA gives rise to a lariat-derived circRNA [112, 113]. All these observations suggest that human cells have developed multiple mechanisms to fine-tune the expression of ATXN2L in dependence on diverse feedbacks, as a highly relevant molecule. While both, ATXN2L and ATXN2, are found among the >300 RNA-binding proteins (RBPs) in the stress granule proteome [56, 114–116], it is unclear so far, with which factors they have close interaction and share functional cooperation. In the human cervical cancer cell line HeLa, preferential interactions within SG were reported between ATXN2L, ATXN2, DDX1, FAM98A, and NUFIP2 [117]. Indeed, the relocalization of ATXN2L to SG depends on PRMT1-mediated arginine methylation [118], and FAM98A and DDX1 are activators of this post-translational modification [119], so FAM98A and DDX1 seem to be upstream factors that determine the trafficking changes of specific ribonucleoproteins (RNPs) during the switch from growth to stress periods. The role of the polysome-associated factor and DDX6 interactor NUFIP2 [120, 121] in this SG subcomplex remains unclear. Another preferential interaction for ATXN2L was claimed with the RNA-binding, cold shock domain-containing protein YBX1 [122, 123]. Variants in the human *ATXN2L* gene have been reported in very few live patients so far [124, 125], who showed developmental delay and macrocephaly.

Here, we performed (i) coimmunoprecipitation and mass spectrometry analyses in WT and ATXN2L-null mouse embryonic fibroblasts (MEF), (ii) global proteome profiling via label-free mass spectrometry in wildtype (WT) and ATXN2L-null MEF, (iii) the analysis of ATXN2L interactor proteins in 14-month-old spinal cords from mice with progressive ATXN2 aggregation due to the *Atxn2*-CAG100-KnockIn (KIN) mutation as established SCA2 model [36, 126–132]. With this strategy, we aimed to identify protein interactors that depend on ATXN2L in their abundance, and undergo changes during the motor neurodegeneration process in SCA2 models. Our work elucidates the pathways that are important for the survival of differentiated neurons under stress.

## Materials and Methods

### MEF culture, treatment, coimmunoprecipitation, and proteome profiling

The generation and culture of 5 WT and 5 ATXN2L-null MEF lines was done as previously reported [106], and their exposure to sodium arsenite (NaARS) stress also followed published protocols [129]. In these experiments, the cells were plated in 175 cm^2^ flasks and allowed to grow until they reached about 90% confluence before being treated with 0.5 mM NaARS (Sigma Aldrich, St. Louis, MO, USA, #S7400) for 45 minutes. Then, MEFs were ruptured in a lysis buffer - 20 mM Tris/HCl pH 8.0, 137 mM NaCl, 2 mM EDTA, 1% NP40, 1% glycerol with Protease-Inhibitor Cocktail cOmplete (Roche Diagnostics, Mannheim, Germany) - via 30 min head-to-head rotation at 4 °C. Nuclear debris was removed via centrifugation at full speed at 4 °C for 20 min. The protein content was determined via BCA (Life Technologies, Karlsruhe, Germany). A total of 1000 μg of protein lysate was incubated with 4 μg of pull antibody (anti-ATXN2L from Invitrogen, Carlsbad, CA, USA #PA5-59601, or normal rabbit IgG from Cell Signaling Technology, Danvers, MA, USA #2729) and rotated for 2 h at room temperature (RT). In the meantime, 1.5 mg of Dynabeads (Thermo Fisher, #10004D) was washed 3 times with PBS/T and added to the lysate/antibody solution. The mix was rotated head-to-head for 60 min at RT, to be either analyzed by immunoblot as described in the following sentences, or by nano LC-MS/MS as described in the subsequent paragraph. For immunoblots, 5 washes with PBS were performed, then the tubes were fixed on a magnetic stand and elution was carried out with 40 μL of 50 mM glycine, pH 2.8. The eluate was mixed with a loading buffer, boiled for 5 min at 90 °C, and loaded for SDS electrophoresis. The antibodies used for co-IP detection were ATXN2L (Proteintech, Rosemont, IL, USA, #24822-1-AP) or PABP (Abcam, Cambridge, UK, #ab21060).

### “On beads” digest for Co-IP samples

Four MEF coimmunoprecipitation samples on magnetic beads (Thermo Scientific Pierce, Waltham, MA, MS-Compatible Magnetic IP Kits, catalogue no: 90409) were washed 3 times in 100 mM ammonium bicarbonate buffer. This was followed by tryptic digestion including reduction and alkylation of cysteines. Reduction was performed by adding tris(2-carboxyethyl)phosphine to a final concentration of 5.5 mM at 37 °C on a rocking platform (600 rpm) for 30 min in a total volume of 150 µL. For alkylation, chloroacetamide was added to a final concentration of 24 mM at room temperature on a rocking platform (600 rpm) for 30 min. Proteins were then digested with 100 ng trypsin (Roche, Basel, Switzerland) per sample at pH 8, shaken at 800 rpm at 37 °C for 18 h. The samples were acidified by adding 3.75 µL of 100% formic acid (2% final concentration), centrifuged briefly and placed on the magnetic rack. The supernatants containing the digested peptides were transferred to a new low-protein binding tube. Peptide desalting was performed on self-packed C18 columns in a tip. Eluates were lyophilized and reconstituted in 38 µL of 5% acetonitrile and 2% formic acid in water, briefly vortexed and sonicated in a water bath for 30 sec before injecting 20 µL onto a nano-LC-MS/MS.

### Global proteomics

10 MEF samples (Atxn2l KO and WT) were lysed under denaturing conditions in 300 µl of a buffer containing 3 M guanidinium chloride (GdmCl), 10 mM tris(2-carboxyethyl)phosphine, 40 mM chloroacetamide, and 100 mM Tris-HCl pH 8.5. Lysates were denatured at 95°C for 10 min shaking at 1000 rpm in a thermal shaker and sonicated in a water bath for 10 min. The protein concentration of each sample was measured with a BCA protein assay kit (23252, Thermo Scientific, USA). 30 µg of each sample was diluted with a dilution buffer containing 10% acetonitrile and 25 mM Tris-HCl, pH 8.0, to reach a 1 M GdmCl concentration. Then, proteins were digested with LysC (Roche, Basel, Switzerland; enzyme to protein ratio 1:50, MS-grade) shaking at 700 rpm at 37°C for 2 hours. The digestion mixture was diluted again with the same dilution buffer to reach 0.5 M GdmCl, followed by tryptic digestion (Roche, enzyme to protein ratio 1:50, MS-grade) and incubation at 37°C overnight in a thermal shaker at 700 rpm. Peptide desalting was performed according to the manufacturer’s instructions (Pierce C18 Tips, Thermo Scientific, Waltham, MA). Desalted peptides were reconstituted in 0.1% formic acid in water and further separated into four fractions by strong cation exchange chromatography (SCX, 3M Purification, Meriden, CT). Eluates were first dried in a SpeedVac, then dissolved in 5% acetonitrile and 2% formic acid in water, briefly vortexed, and sonicated in a water bath for 30 seconds prior injection to nano-LC-MS/MS.

### LC-MS/MS instrument settings for shotgun proteome profiling and data analysis

Liquid chromatography–tandem mass spectrometry (LC-MS/MS) was performed using nanoflow reversed-phase liquid chromatography (Dionex Ultimate 3000, Thermo Scientific) coupled online to a Q-Exactive HF Orbitrap mass spectrometer (Thermo Scientific), as previously reported [133]. Briefly, LC separation was performed using a PicoFrit analytical column (75 μm ID × 50 cm long, 15 µm tip ID; New Objectives, Woburn, MA) packed in-house with 3 µm C18 resin (Reprosil-AQ Pur, Dr Maisch, Ammerbuch, Germany). Peptides were eluted using a gradient of 3.8 to 38% solvent B in solvent A over 120 min at a flow rate of 266 nL per minute. Solvent A was 0.1% formic acid and solvent B was 79.9% acetonitrile, 20% H2O and 0.1% formic acid. The nanoelectrospray was generated by applying 3.5 kV. A cycle of one full Fourier transform scan mass spectrum (300-1750 m/z, resolution of 60,000 at m/z 200, automatic gain control (AGC) target 1 × 106) was followed by 12 data-dependent MS/MS scans (resolution of 30,000, AGC target 5 × 105) with a normalized collision energy of 25 eV. A dynamic exclusion window of 30 sec was used to avoid multiple sequencing of the same peptides.

Raw MS data were processed using MaxQuant software (v2.2.0.0) and searched against the *Mus musculus* proteome database UniProtKB UP000000589 with 55,260 protein entries, released in June 2023. MaxQuant database search parameters were a false discovery rate (FDR) of 0.01 for proteins and peptides, a minimum peptide length of seven amino acids, a first search mass tolerance for peptides of 20 ppm, and a main search tolerance of 4.5 ppm. A maximum of two missed cleavages was allowed for tryptic digestion. Cysteine carbamidomethylation was set as a fixed modification, while N-terminal acetylation and methionine oxidation were set as variable modifications. The MaxQuant processed output files can be found in Supplementary Table 1, showing peptide and protein identification, accession numbers, % sequence coverage of the protein and q values. The mass spectrometry data have been deposited at the ProteomeXchange Consortium (http://proteomecentral.proteomexchange.org) via the PRIDE partner repository [134] with the data set identifier PXD061048 for MEF proteome profiles, and PXD061161 for MEF Co-IPs.

### Fluorescent immunocytochemistry

The microscopic detection of ATXN2L or ATXN2 compared with NUFIP2 in MEF cultures, without versus with NaARS-mediated oxidative stress to validate their co-localization in stress granules, was performed as previously described [129], using the following primary antibodies: ATXN2L (Proteintech, #24822-1-AP) with NUFIP2 (Proteintech, #67195-1-Ig), ATXN2 (BD Transduction, #611378) with NUFIP2 (Proteintech, 17752-1-AP).

### Breeding, aging, genotyping, and dissection of *Atxn2*-CAG100-KnockIn mice

Spinal cord tissues from 14-month-old mice were obtained as previously described [129, 131] and stored at -80 °C until analysis. The study was ethically assessed by the Regierungspraesidium Darmstadt, with approval number V54-19c20/15-FK/1083.

### Spinal cord protein extraction and quantitative immunoblotting

Mouse cervicothoracic spinal cord tissues were homogenized with a motor pestle in 5-10x weight/volume amount of RIPA buffer consisting of 50 mM Tris-HCl (pH 8.0), 150 mM NaCl, 2 mM EDTA, 1% Igepal CA-630 (Sigma Aldrich, St. Louis, MO), 0.5% sodium deoxycholate, 0.1% SDS, cOmplete™ Protease Inhibitor Cocktail (Roche), and Halt™ Phosphatase Inhibitor Cocktail (Thermo Fisher Scientific). Similarly, MEF and human skin fibroblast pellets were homogenized in RIPA buffer by pipetting. The resulting protein suspensions were sonicated, and protein concentration was determined in a Tecan Spark plate reader (Tecan Group Ltd, Männedorf, Switzerland) using a Pierce™ BCA protein assay kit (Thermo Fisher Scientific). 15 to 25 μg of total proteins were mixed with 6x loading buffer consisting of 250 mM Tris-HCl pH7.4, 20% glycerol, 4% SDS, 10% 2-mercaptoethanol, and 0.005% bromophenol blue, incubated at 90 °C for 5 min, separated on 8-15% polyacrylamide gels at 120 Volts, and transferred to nitrocellulose membranes (0.2 μm) (Bio-Rad Laboratories, Hercules, CA). The nitrocellulose membranes were blocked in 5% BSA/TBS-T, and incubated overnight at 4 °C with primary antibodies. Afterwards, the nitrocellulose membranes were incubated for 1 h at room temperature, with fluorescently labeled secondary IRDye® 800CW goat anti-mouse (LI-COR 926-32210, 1:10,000), IRDye® 800CW goat anti-rabbit (LI-COR 926-32211, 1:10,000), IRDye® 680RD goat anti-mouse (LI-COR 926-68070, 1:10,000) or IRDye® 680RD goat anti-rabbit (LI-COR 926-68071, 1:10,000). Membranes were scanned using an Odyssey® Classic Imager. Image visualization and quantification of signal intensities were performed using Image StudioTM software (version 5.2) (LI-COR Biosciences, Lincoln, NE). The following primary antibodies were used: ATXN2L (Proteintech, Rosemont, IL, USA, #24822-1-AP), ATXN2 (Proteintech, #21776-1-AP), NUFIP2 (Proteintech, #17752-1-AP), NUFIP1 (Proteintech, #12515-1-AP), SYNE2 (Invitrogen, #PA5-78438) and DCLK1 (Cell Signaling Technology, #62257). GAPDH (Calbiochem, San Diego, CA, USA, #CB1001, 1:10000) served as loading controls. Precision Plus Protein™ All Blue Prestained Protein Standards (Bio-Rad, Hercules, CA, USA, #1610373) was used as size marker.

### Reverse transcriptase quantitative real-time polymerase chain reaction (RT-qPCR)

Total RNA isolation from aged spinal cord tissue and MEF pellets was performed with TRIzol Reagent (Sigma Aldrich) according to manufacturer’s instructions. Total RNA yield and purity were quantified using a Tecan Spark plate reader at 230, 260, and 280 nm, in a NanoQuant plate. cDNA synthesis was performed from 1 μg of total RNA template using the SuperScript IV VILO kit (Invitrogen) according to the manufacturer’s instructions. Gene expression profiles were assessed by RT-qPCR using a StepOnePlusTM (96 well) Real-Time PCR System (Applied Biosystems, Foster City, CA, USA). RT-qPCRs were run in technical duplicates on cDNA from 25 ng total RNA, with 1 μl TaqMan® Assay, 10 μl FastStart Universal Probe Master 2× (Rox) Mix (Roche) and ddH2O up to 20 μl of total volume. The PCR cycling conditions were 50 °C for 2 min, 95 °C for 10 min, followed by 40 cycles of 95 °C for 15 min and 60 °C for 1 min. The gene expression TaqMan® assays (Thermo Fisher Scientific, Waltham, Massachusetts, USA) used for this study were: *Atxn2* (Mm01199894_m1), *Atxn2l* (Mm00805548_m1), *Cux1* (Mm01195598_m1), *Dclk1* (Mm01545304_m1), *Nufip1* (Mm00479126_m1), *Nufip2* (Mm01077988_m1), *Syne1* (Mm04238399_m1), *and Syne2* (Mm00621101_m1). The data were analyzed via the 2^−ΔΔCt^ method [135], using *Tbp* (Mm00446973_m1) as housekeeping transcript.

### Statistics and graphical presentation

Unpaired Student t-tests with Welch’s correction were used to establish comparisons for continuous variables between homozygous *Atxn2*-CAG100-KIN and WT animals. Bar charts depicting the mean and standard error of the mean (SEM) values were used for data visualization. GraphPad (Version 10.4.1, for Windows, GraphPad Prism, Boston, MA) software was used for all statistical analyses and Volcano plot generation. Significance was assumed at p<0.05 and highlighted with asterisks: *p<0.05, **p<0.01, ***p<0.001.

## Results

### Coimmunoprecipitation of ATXN2L interactors in WT versus ATXN2L-null MEFs shows novel interactions, stronger than the known LSM12, ATXN2, PABPC1 and G3BP2 associations

To identify the strongest ATXN2L interactors inside and outside RNA granules, we used previously described MEF derived from mice with constitutive deletion of *Atxn2l* exons 5-8 [106]. Co-IP pulling was done with a polyclonal antibody against human ATXN2L residues 456-547 (Invitrogen, PA5-59601) or with unspecific immunoglobulin-G as negative control (IgG control, Cell Signaling Technology, #2729S). Comparisons were performed of unstressed WT MEF versus NaARS-stressed WT MEF, and of WT versus homozygous ATXN2L-null MEF. Preliminary immunoblot analysis of these samples (1000 µg each) with antibodies against human ATXN2L residues 712-Cterm (Proteintech, 24822-1-AP) demonstrated the expected presence of ATXN2L with its known interactor PABPC1 (Figure 1).

**Figure 1.**
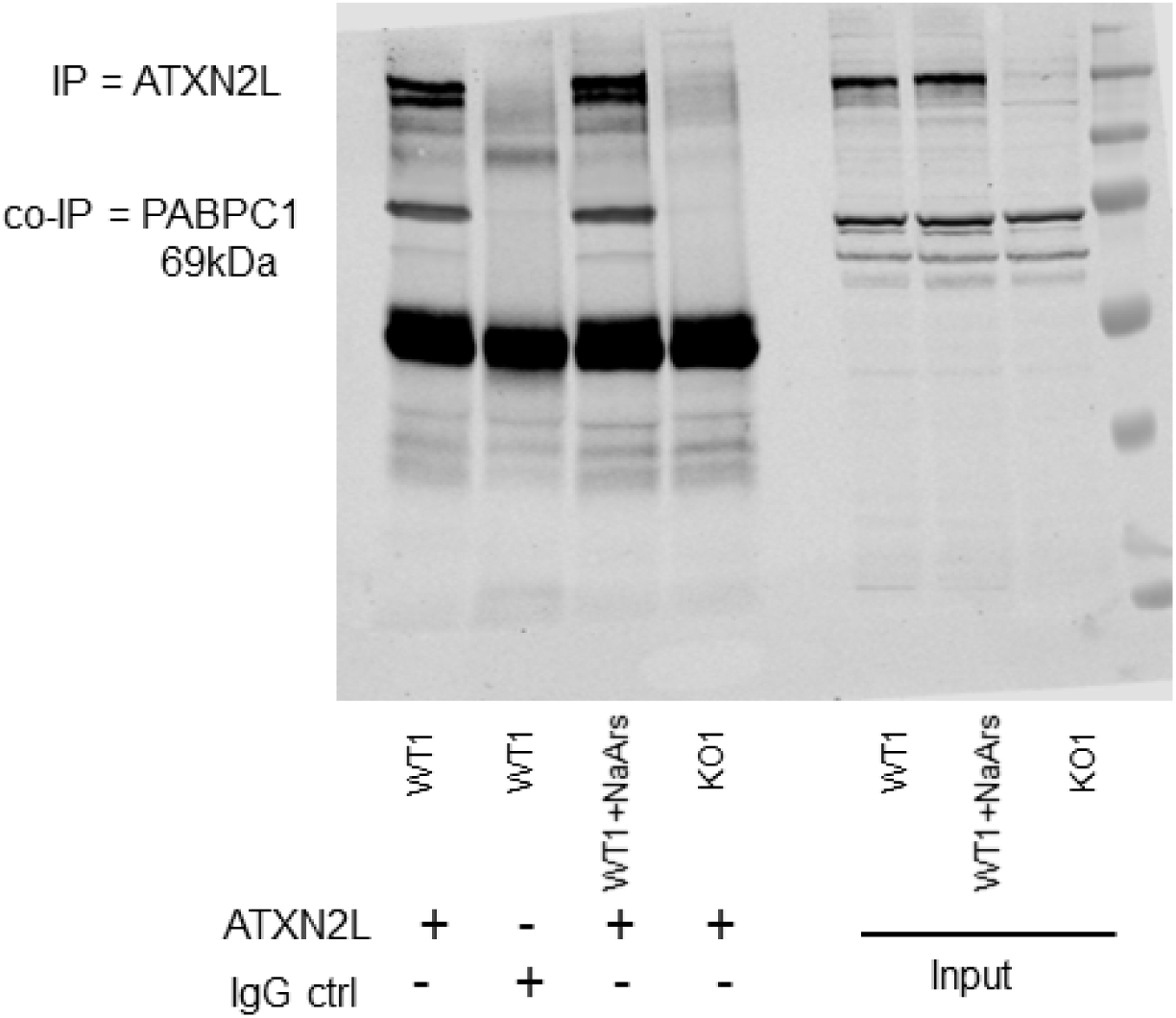
Immunoblot analysis of ATXN2L co-IP samples. Co-IP samples were pulled with an antibody against ATXN2L or with unspecific immunoglobulin-G as negative control (IgG). Comparisons were performed of unstressed WT MEF versus NaARS-stressed WT MEF, and of WT versus homozygous ATXN2L-null MEF. Immunoblot analysis of these samples (1000 µg each) with antibodies ATXN2L demonstrated the expected presence of ATXN2L with its known interactor PABPC1.

After identifying and quantifying the polypeptide components of these coimmunoprecipitates via LC-MS/MS and MaxQuant software (Table S1), 29 proteins were found associated with ATXN2L in unstressed WT, but neither in IgG ctrl nor ATXN2L-null samples (Table 1), as the most credible candidates. In comparison with the known ATXN2L cytosolic interactors and SG components ATXN2, PABPC1 and G3BP2, 24 proteins had higher abundances that approached the high concentration of ATXN2L (Table 1) and were therefore more likely candidates for equimolar binding. Prominently, the levels of label-free quantification (LFQ) as approximate reflection of protein abundance identified RNA-binding proteins (RBP) PABPN1, NUFIP2, MCRIP2, RBMS1, LARP1, PTBP1, FMR1, RPS20, FUBP3, MBNL2, ZMAT3, SFPQ, CSDE1, HNRNPK and HNRNPDL, to show association with ATXN2L at approximately equimolar ratios, with more or similar strength relative to known interactors such as LSM12, ATXN2, PABPC1, G3BP2, being completely absent from the KO co-IP. The only non-RNP with this pattern were the actin complex components SYNE2, LMOD1, ACTA2, FYB and GOLGA3, the nuclear histone HIST1H1C, the pore-forming immunity factor MPEG1, and the peroxidase PRDX1. Clearly, there is an enrichment of RBP and actin cytoskeleton factors, and this is in excellent agreement with published *Drosophila melanogaster* data implicating dATX2 in polysomal translation and in actin filaments [34, 51]. In the discussion below, the detailed comparison of ATXN2L interactor candidates with relevant literature confirmed the credibility and relevance of the mass spectrometry data. Thus, these novel observations will deepen our understanding of ATXN2L functions at the molecular level, and further validation experiments were done to define which of these proteins depend on ATXN2L regarding their stability.

**Table 1.**
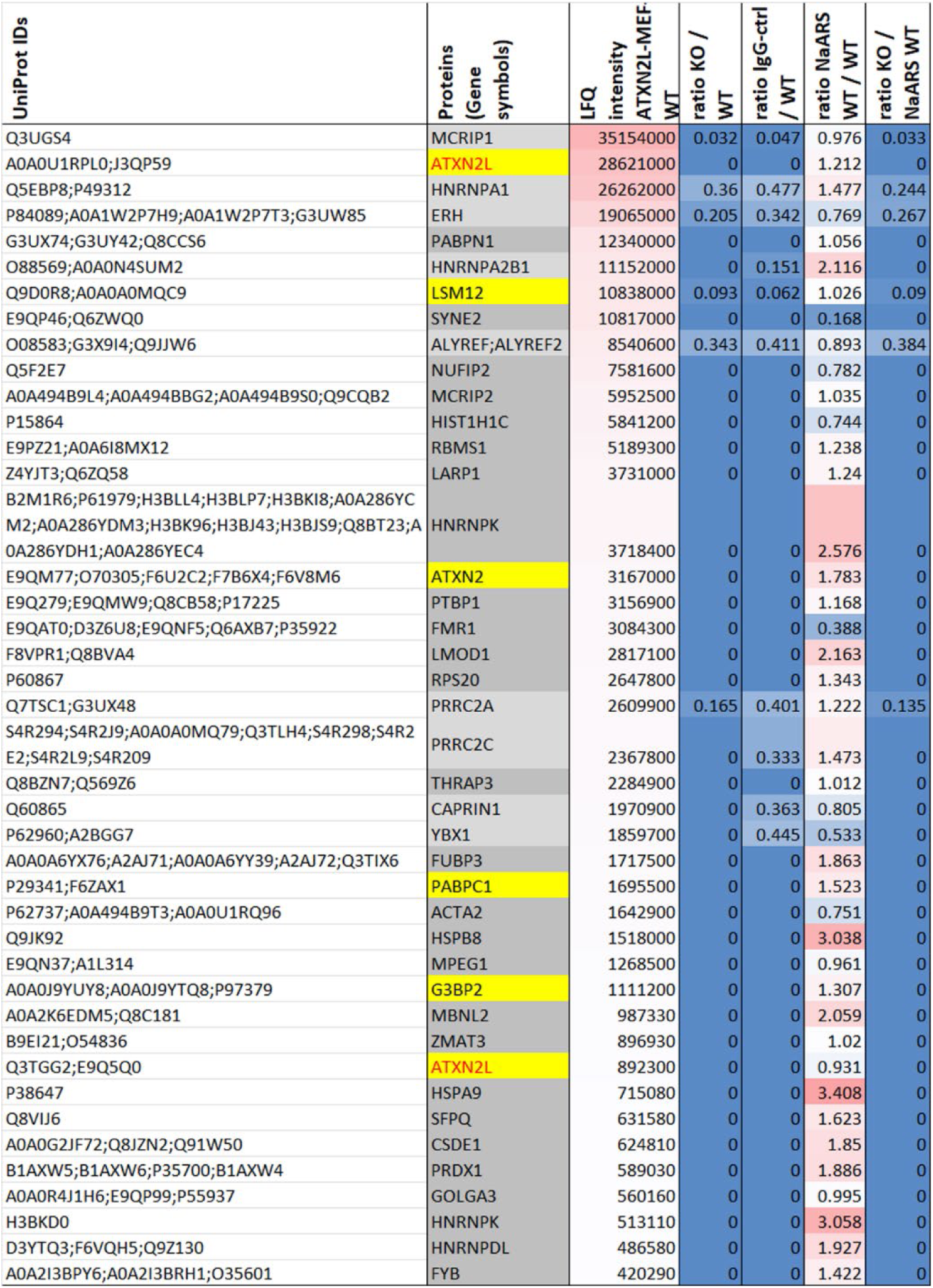
Mass spectrometry quantifications. Mass spectrometry identification and quantification of ATXN2L co-IP components in MEF, without stress or after NaARS stress. ATXN2L and its known interactor proteins are highlighted in yellow cells; novel interactors that are absent from controls in dark grey; potential interactors with weaker ratios in light grey; protein abundance is illustrated by red background for high values and ratios above 1, while blue background illustrate ratios below 1; very low ratios are shown in dark blue.

MEF exposure to oxidative stress by administration of NaARS triggered strong increases of the protein chaperones HSPB8 / HSPA9 and the RNA chaperone HNRNPK (>2.5-fold) in the ATXN2L co-IP, in parallel to a strong decrease of SYNE2 amounts (<0.25-fold). These observations may simply reflect the reduction of RNP granules that are associated with the cytoskeletal trafficking machinery, with the concurrent induction of refolding efforts in stress granules under conditions of liquid-liquid phase separation, after cell damage.

Strong colocalization of ATXN2L and ATXN2 with NUFIP2 in stress granules induced by NaARS administration in MEF cultures was confirmed in triple immunofluorescence microscopy (Fig. S1 and S2). Thus, both biochemical and morphological observations are compatible with the notion that ATXN2L and NUFIP2 might cooperate functionally.

### Global proteome profiling of MEF reveals NUFIP2 and SYNE2 depletion as consequence of ATXN2L absence

To document ATXN2L-null triggered downstream effects on steady state protein levels across the global proteome, we used the MEF cells again [106]. This unbiased survey of 5 ATXN2L-null versus 5 WT MEF lines by label free mass spectrometry achieved detection of 4386 proteins, including 280 factors that showed upregulation with nominal significance versus 292 factors with downregulation (Table S2). The most credible subset of dysregulated proteins with actual significance and at least two-fold change is shown in a volcano plot (Figure 2). As the main findings, the absence of ATXN2L causes deficiency of two of its interactors; firstly, the reduction of ATXN2L peptides to 8% caused a similar decrease to 8% for NUFIP2 (Nuclear Fragile X Mental Retardation Protein Interacting Protein 2), as DDX6-binding protein with SG localization [117, 136]. Secondly, a reduction to 12% was observed for SYNE2 peptides (also known as Nesprin-2) as actin cytoskeleton movement factor [137–139] that is paralogous to SYNE1 as disease gene responsible for the autosomal recessive ataxia SCAR8 [140, 141]. Otherwise, a very large and significant downregulation to 11% was found for CUX1, as a transcription factor that identifies pyramidal neurons from cortex layers II/III [142–148]. The biggest upregulation to 720% concerned the microtubule polymerization regulator DCLK1 that is responsible for retrograde transport and neuronal migration [149–151].

**Figure 2.**
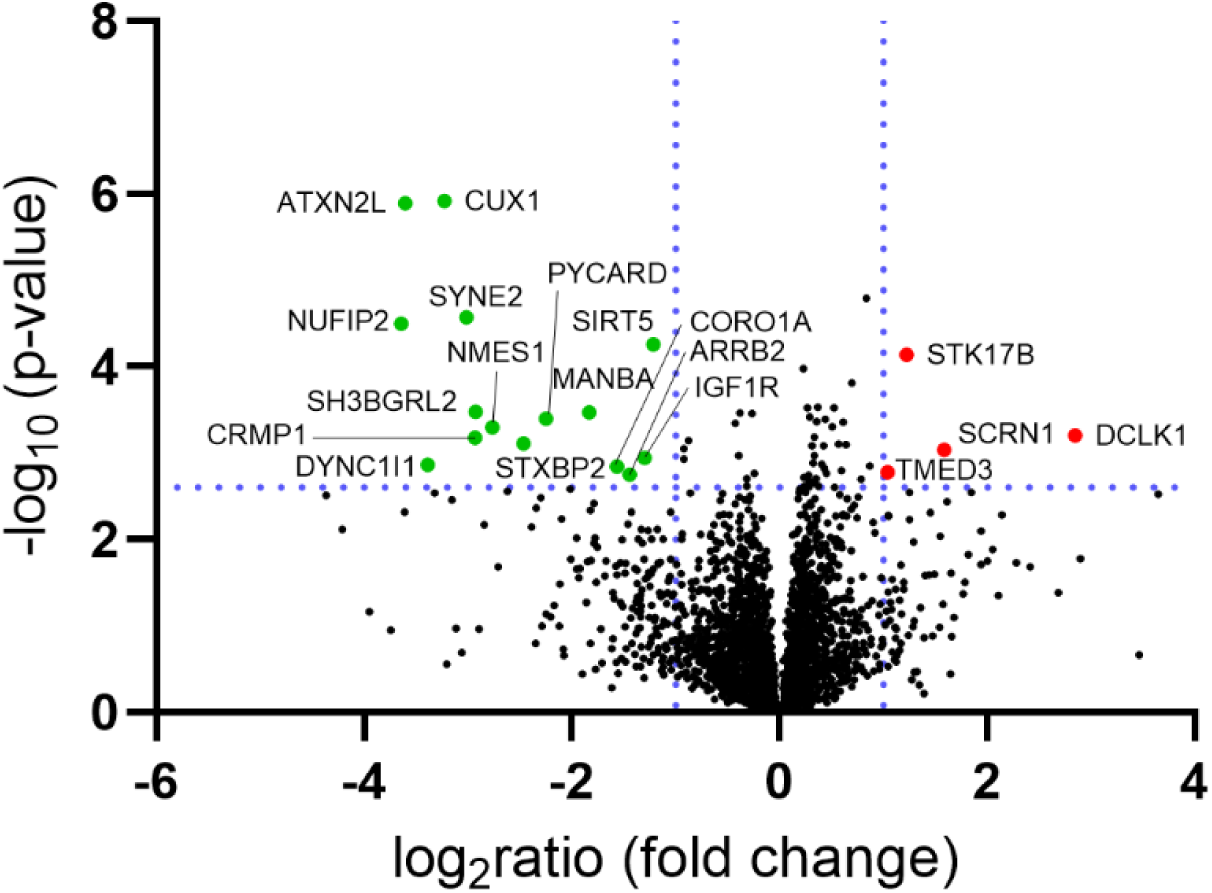
Volcano plot of global proteome profile of ATXN2L-null MEF cells. ATXN2L and its interactor proteins NUFIP2 / SYNE2 as well as pyramidal neuron marker CUX1 are similarly depleted, and several cytoskeletal trafficking pathway components (e.g. DYNC1I1, CRMP1, SH3BGRL2, STXBP2, and DCLK1) appear dysregulated. The position of downregulated factors on the left side was shown as green dots, upregulated factors on the right side as red dots.

Altogether, the loss of the RBP ATXN2L in MEF destabilized NUFIP2 exclusively among its many RBP interactors, and SYNE2 prominently among the actin filament modulators. While ATXN2L did not appear essential for the other RNPs, its loss triggered a spectrum of downstream cytoskeleton dysregulations that range from actin bundling to microtubular transport, involving also vesicle dynamics.

### Validation experiments by quantitative immunoblots and RT-qPCR confirm ATXN2L-null MEFs to show depleted NUFIP2/NUFIP1 protein, reduced SYNE2 protein and *Cux1* mRNA, versus increased DCLK1 expression

Attempting to assess the validity of the principal findings, first the commercially available antibodies were used in quantitative immunoblots (Figure 3A) to demonstrate that two isoforms of ATXN2L near 150 kDa molecular weight were absent in mutant MEFs, while its less abundant paralog ATXN2 showed significantly elevated levels. NUFIP2 had a depletion to 10.6%, its paralog NUFIP1 was reduced to 40.2% and SYNE2 to 61.5%, whereas DCLK1 abundance exhibited a 5-fold elevation. Antibodies against the paralog SYNE1 and against CUX1 were not sensitive or specific enough to generate convincing immunoreactivity in the MEFs.

**Figure 3.**
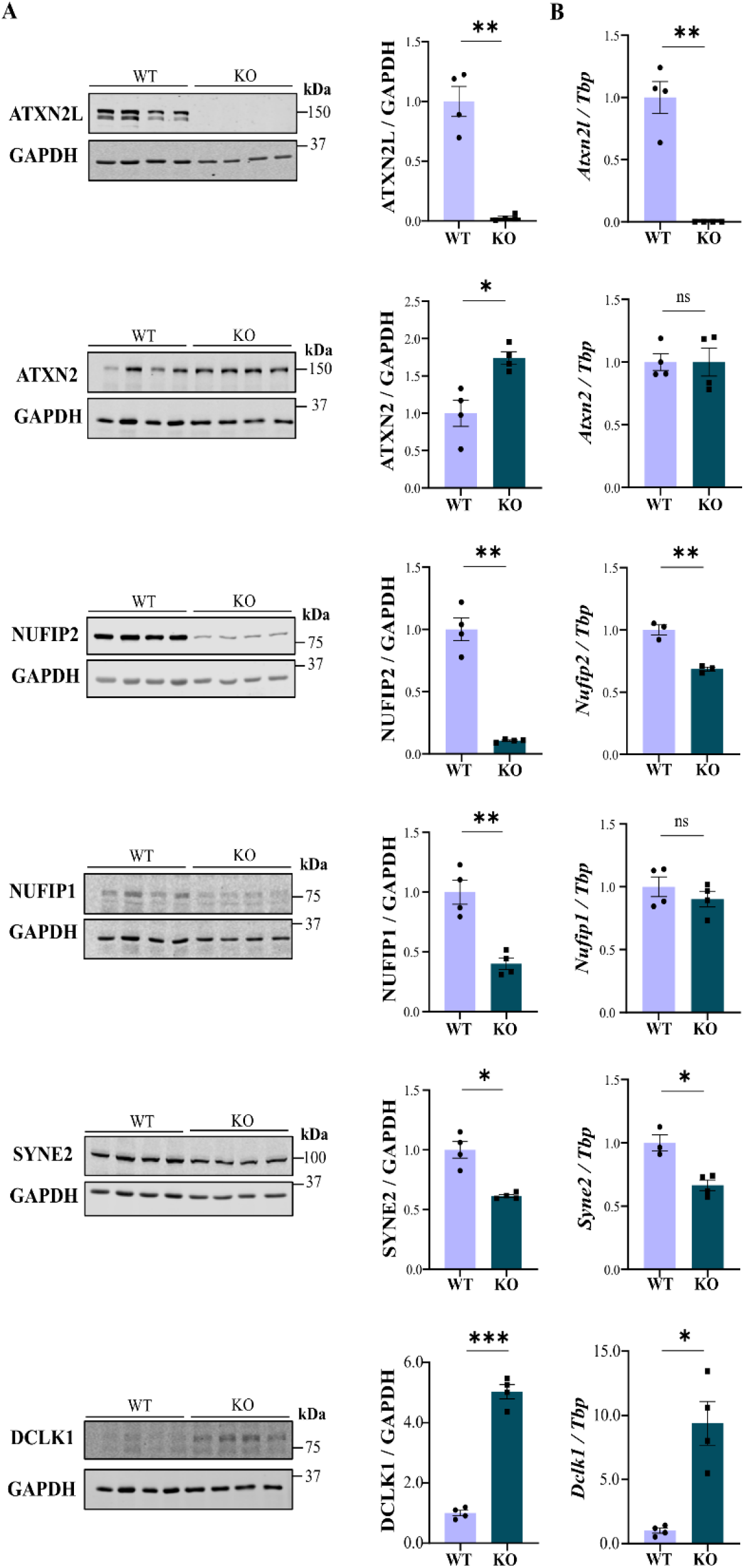
Validation experiments in ATXN2L-null MEFs. **(A)** Quantitative immunoblots document the abundances of proteins of interest in 4 mutant versus 4 WT samples, normalized to GAPDH as loading control. **(B)** RT-qPCR quantifies expression of transcripts of interest in 4 mutant versus 4 WT samples, normalized to *Tbp* mRNA as loading control. Significance was assumed at p<0.05 and highlighted with asterisks: *p<0.05, **p<0.01, ***p<0.001, ns = non-significant.

Employing the RT-qPCR technique to test mRNA expression levels of these factors, the absence of *Atxn2l* transcript was confirmed but no upregulation of its paralog *Atxn2* was confirmed (Figure 3B), suggesting that post-transcriptional mechanisms have to be responsible for the accumulation of ATXN2 protein in ATXN2L-null MEFs. Interestingly, the depletion of NUFIP2 protein was not counteracted by its transcriptional induction, but instead accompanied by reduced *Nufip2* mRNA, suggesting that the loss-of-function of NUFIP2 is beneficial for ATXN2L-null cells. Similarly, the reduced expression of *Syne2* and *Syne1* as well as *Cux1* paralleled their lowered protein abundance. Conversely, a strong increase of *Dclk1* transcript levels was observed to underlie the elevated DCLK1 protein abundance.

Overall, these data reproduce key findings of the proteome profile and point to a beneficial compensatory role of these dysregulations in ATXN2L-null MEFs.

### The ribonucleoprotein aggregation process in the spinal cord of 14-month-old *Atxn2*-CAG100-KnockIn mice includes the accumulation of ATXNL2 and NUFIP2 proteins

We reasoned that the above findings may be relevant for SCA2 disease mechanisms, in view of the known heteromultimerization of ATXN2 with ATXN2L in neuronal RNA granules during transport, and in SG. The polyQ expansion within ATXN2 leads to a ribonucleoprotein aggregation process that sequestrates ribonucleoprotein interactors like PABPC1 and TDP-43 into cytosolic inclusion bodies within spinal motor neurons, possibly also within cerebellar Purkinje neurons [9, 36, 131]. Upon testing ATXN2L with its main protein interactors in mice with ATXN2 polyQ expansion, aged spinal cord tissues showed the previously reported reduction in translated soluble ATXN2 (Figure 4). The well-established ATXN2 aggregation process in aged spinal cord of these KIN mice [131] resulted in significant accumulation of ATXN2L (1.53-fold, p=0.0052) similar to TDP-43 (1.57-fold, p=0.003), together with NUFIP2 monomers (1.56-fold, p=0.021) and homodimers (3.28-fold, p=0.0007), SYNE1 (1.54-fold, p=0.011; SYNE2 analysis did not produce convincing immunoreactivity in these tissues, in agreement with GeneCards database proteome entries that SYNE1 but not SYNE2 is detected by mass spectrometry in spinal cord) and DCLK1 (1.25-fold, p=0.048). The sequestration of these factors into the phase-separated inclusion bodies would reduce their availability in the water-soluble cytosolic fractions. Thus, a partial loss-of-function due to aggregation or a gain-of-function due to excess amounts of these ATXN2L interactors might contribute to the cellular pathology in SCA2 via altered RNA processing and cytoskeletal dynamics.

**Figure 4.**
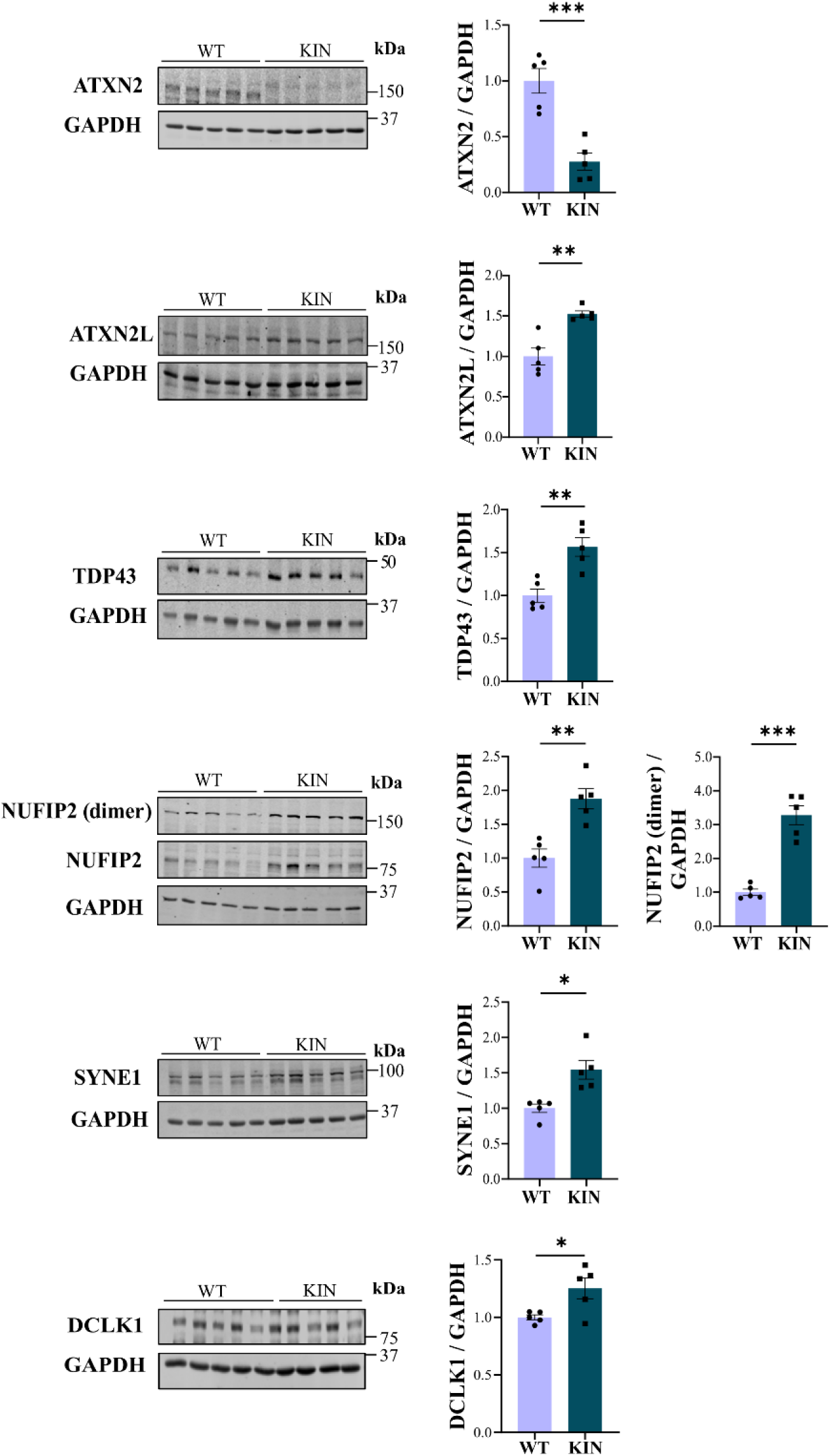
Validation experiments in the spinal cord of end-stage *Atxn2*-CAG100-KnockIn mice. Quantitative immunoblots demonstrating the deficiency of polyQ-expanded ATXN2 in the soluble compartment, together with accumulation of TDP-43, ATXN2L, NUFIP2 (monomers at 75 kDa, homodimers at 150 kDa), SYNE1 and DCLK1 in spinal cord homogenates from 14-month-old *Atxn2*-CAG100-KnockIn mice (graphs reflect 5 mutant versus 5 WT samples), using GAPDH as loading normalizer. Significance was assumed at p<0.05 and highlighted with asterisks: *p<0.05, **p<0.01, ***p<0.001.

## Discussion

ATXN2L is essential for embryonic development in mice, indicating its crucial role in cellular survival. However, there is very limited knowledge about this protein based on patient genetics, biochemistry, microscopy studies, and bioinformatics predictions. Previously, it was thought that ATXN2L and ATXN2 function within ribonucleoprotein complexes, which are relatively stationary in the spliceosome or near the rough endoplasmic reticulum. They are believed to modulate the ribosomal translation pathway during cellular growth phases and assist in RNA quality control pathways during cellular stress and repair phases.

New evidence from this study indicates two important findings regarding ATXN2L in mice. First, the association of ATXN2L with various ribonucleoproteins is essential for NUFIP2. Second, ATXN2L interacts with SYNE2 and several other components of the actin-microtubule cytoskeleton, suggesting that it plays a role in RNA surveillance during polarized transport from the nuclear spliceosome to the tips of cell processes.

In the sections below, we will revisit our experimental data in detail and integrate it with the current literature to propose a credible scenario regarding the functions of ATXN2L.

### Coimmunoprecipitation findings

In agreement with previous observations that ATXN2L is not exclusively cytosolic like ATXN2 [100], but instead shows prominent nuclear localization [118], the ATXN2L coimmunoprecipitates contained not so much the cytosolic poly(A) binding protein PABPC1, but mainly the nuclear poly(A) binding protein PABPN1 (in control of nuclear exosomal degradation of polyadenylated RNAs) [152–156], with >7-fold higher LFQ intensity as measure of abundance. Furthermore, the ATXN2L co-IP contained ZMAT3 (also known as Wig-1), which localizes mainly to the nucleolus, binding to long double-strand RNA (dsRNA) and short microRNA-like dsRNA to regulate splicing and cell senescence, while shuttling to the cytosol to regulate ARE-mediated RNA decay there [157–161]. In addition, the ATXN2L co-IP also comprised a subcomplex consisting of three nuclear speckle-paraspeckle proteins that modulate selective expression and translation of internal ribosomal entry site (IRES) sequences [162]: (i) THRAP3 (also known as TRAP150), an RNA alternative splicing and nuclear RNA decay mediator, as well as R-loop resolution factor [163, 164]; (ii) SFPQ (also known as PSF or Polypyrimidine Tract-Binding Protein-Associated-Splicing Factor that contains a K-homology domain) as known interactor of THRAP3 and of ALS-associated NEAT1 lncRNA [165–170]; (iii) PTBP1 (also known as HNRNPI or as Polypyrimidine Tract Binding Protein 1 [171, 172]. Providing strong evidence for the nuclear localization of ATXN2L, the co-IP included the nuclear linker histone HIST1H1C (which is also known as H1.2, in control of cell growth versus apoptosis following DNA double-strand breaks, [173–175]). These observations implicate ATXN2L in nuclear RNA splicing and degradation, as well as dsRNA processing, for the adaptation of cells to stress conditions.

The highest number of ATXN2L interactors were RBPs. Those with predominant localization in the nucleus included MCRIP2 (also known as FAM195A, a DDX6 interactor and SG component [121]), HNRNPK (binding to polyC sequences [176, 177]), FUBP3 (also known as MARTA2, binding 3’ untranslated region UAU sequences, containing a K-homology domain [178–180]), MBNL2 (as RNA alternative splicing and localization mediator, which contains a K-homology domain, binding to the ACACCC zipcode core sequence and to triplet repeat structures [181–183]) and HNRNPDL (also known as AU-Rich Element RNA-Binding Factor or JKTBP, [184–187]). Those with distribution throughout nucleus and cytoplasm were firstly FMR1 (Fragile X Messenger Ribonucleoprotein 1) as mRNA nuclear export / alternative splicing / dendritic transport factor that contains a K-homology domain and binds polyU sequences, again with known SG relocalization [188, 189]; secondly NUFIP2 (Nuclear FMR1 Interacting Protein 2, another DDX6 interactor and SG component [121]); thirdly LARP1 (La Ribonucleoprotein 1 Translational Regulator, again in the DDX6 interactome and SG component, [189–191]; and fourth RPS20 (a component of the small ribosome subunit for mRNA quality control, which acts in nucleolus to control cell proliferation [192–194]). Mutation of mouse ATXN2 was previously shown to affect ribosomal protein abundance [60], but the very selective ATXN2L interaction with RPS20 is noteworthy. Binding between Ataxin-2 and DDX6 is well established from human to flies [27], so also the enrichment of DDX6 interacting proteins in the ATXN2L co-IP is expected. Predominant cytoplasmic distribution and known SG relocalization was represented only by G3BP2 [195–197]. G3BP2 and G3BP1 are distinct from other SG proteins, since they contain SH3 domains that are expected to interact with proline-rich motifs (PRM), and indeed ATXN2L and ATXN2 homologs across phylogenesis contain at least two conserved functional PRMs [37, 198, 199].

The LSm and LSm-AD motifs of ATXN2L act as RNA chaperone that corrects conformation without consuming ATP energy, and may interact with DDX6 as RNA helicase that corrects conformation more forcefully with energy from ATP. Therefore, it seems logical that the ATXN2L interactome is enriched in other RNA chaperones such as HNRNPI, HNRNPK, HNRNPDL [200], together with additional factors that contain a K-homology domain. Already in bacteria, a cooperation between the Sm domain containing factor Hfq and the K homology domain containing factor PNPase (polyribonucleotide nucleotidyltransferase) was found crucial for riboregulation [201].

Overall, the ATXN2L co-IP findings above are credible and indicate that the main role of ATXN2L is focused on RNA surveillance, with preference to the nucleus.

In view of the MBNL2 role for RNA relocalization along the cytoskeleton, it was interesting to note that the RNA-binding protein RBMS1 appeared together with cytoskeletal ACTA2 (actin alpha 2) in the ATXN2L co-IPs, with the single-strand [AU]CU[AU][AU]U-sequence binding RBMS1 known to control ACTA2 transcripts, and to influence differentiation as well as radial migration of neural progenitors [202–204]. The co-IP presence of LMOD1 as actin-nucleating factor [205–207] also suggests a cytoskeleton association. The putative ATXN2L interactor SYNE2 (also known as Nesprin-2) is a component of the LINC (Linker of Nucleoskeleton and Cytoskeleton) complex that regulates cell polarity and nuclear movement by association with dynein-dynactin, kinesin and myosin motor proteins [137, 139, 208–214]. Therefore, SYNE2 association represents additional evidence that ATXN2L localizes near the nuclear envelope along the cytoskeleton. Furthermore, FYB (also known as ADAP) as potential ATXN2L interactor has a role in actin rearrangement [215–217]. In another subcellular context that agrees with the reported Golgi localization of dATX-2 in *Drosophila melanogaster* [218], GOLGA3 as a candidate ATXN2L interactor is implicated in the cell reorientation by microtubules to a position where membrane and glycoprotein delivery for neurite extension is optimized. Its mutations lead to primary ciliary dyskinesia [219–221]. Overall, several putative ATXN2L interactors suggest its involvement in trafficking along the cytoskeleton, presumably together with RNA granules.

The observation of molecular chaperones is frequent in any co-IP experiment, so cytosolic HSPB8 and membrane-associated HSPA9 as heat-shock proteins need not represent a specific ATXN2L function. However, CSDE1 (also known as UNR) as Golgi-/vesicle-associated cold shock protein for RNA stem-loop binding, polypyrimidine-tract-mediated IRES-dependent translation initiation, stress granule assembly and RNA decay appears to be a specific ATXN2L interactor within the RNA surveillance pathway [222–227].

The ATXN2L associated PRDX1 (Peroxiredoxin 1) is an antioxidant defence factor, which would modulate SG assembly/disassembly [228, 229].

In contrast, the identification of MPEG1 (also known as Perforin-2) in the ATXN2L interactome is unrelated to previously mentioned pathways, in view of MPEG1 localization to cytoplasmic vesicles / phagosomes / lysosomes where it forms pores to mediate bacterial destruction and the cytosolic escape of bacterial fragments [230–232]. Thus, the presence of MPEG1 in the ATXN2L co-IP may represent an artefact, or may be part of stress-triggered innate immune responses to toxic dsRNA or malformed RNA.

### ATXN2L-null MEF proteome dysregulations

As the main finding of this paper, NUFIP2 appeared destabilized by the loss of ATXN2L in MEF, and this mass spectrometry observation was validated in independent immunoblot experiments. This massive and selective impact was unexpected, because most ribonucleoprotein interactions within SG are promiscuous, mediated by intrinsically disordered domains (IDR) of proteins and the associated RNAs in liquid-liquid phase separation (LLPS). Such an insight about a quite exclusive interaction points to a specific function that is performed by ATXN2L together with NUFIP2 and support from their binding partner DDX6. Again, not too much has been elucidated about the physiological roles of NUFIP2, since its initial discovery as a nuclear and cytoplasmic shuttling factor with ribosomal function within the FMR1 Fragile X protein complex with RNAs [120, 189], and its subsequent identification as a principal component in the DDX6 protein interactome [121]. On the one hand, NUFIP2 was observed to bind mRNAs within their 3’ untranslated region [233], and similarly Ataxin-2 was also shown to target the 3’UTR [30], in particular AU-rich elements (ARE) that are frequent in growth-regulating mRNAs [234]. On the other hand, NUFIP2 is recruited to lysosomes (together with G3BP1 and GABARAPs of the mATG8 family) where it inactivates MTOR [136, 235], and again Ataxin-2 in yeast, worms and mice was reported to repress MTOR [49, 61, 236–239]. Thus, current literature confirms that both NUFIP2 and ATXN2L are inhibitors of growth signals, and both mediate this effect by binding to specific sequences shortly upstream the poly(A) tail of mRNAs.

Except for NUFIP2 depletion, the other proteome dysregulations did not include RNPs. Instead, they affected multiple cytoskeletal transport factors, among which the actin cytoskeleton movement factor SYNE2 plays the key role as ATXN2L interactor. SYNE2 was depleted in parallel to ATXN2L, and again this mass spectrometry observation was validated by immunoblot experiments. SYNE2 (aka Nesprin-2) is crucial for the tethering of nuclei and their relocation within polarized cells [139, 212–214, 240–244], so its depletion possibly underlies the increased presence of giant multinucleated cells in the ATXN2L-null MEF lines [106]. Therefore, SYNE2 is key for the migration and differentiation of post-mitotic cortical neurons [245], and this indirect effect may contribute to our previous report that ATXN2L-null mice die *in utero* with a phenotype of deficient cortex lamination and neuronal apoptosis [106]. In this context, the downregulation of the nuclear transcription factor CUX1, which is selectively expressed in cortical layer II/III neurons may also contribute to the embryonic lethality of ATXN2L-null mice and perhaps to the motor neuron degeneration during postnatal life of SCA2 patients [142–148]. Although a specific and sensitive antibody to detect CUX1 was not available to us, its decreased mRNA expression levels could be demonstrated in validation experiments.

Downstream indirect effects in the proteome, which may represent pathology effectors or act in compensatory manner, include dysregulations of the microtubule-binding growth cone collapse factor CRMP1 [246], the actin/spectrin interactor and migration modulator SH3BGRL2 [247, 248], the RNA granule retrograde motor DYNC1i1 [249, 250], the actin/vesicle dynamics modulator STXBP2 (also known as MUNC18B) [251], the cytoskeleton-associated membrane protrusion factor CORO1A [252], the cofilin scaffold ARRB2 [253], the myosin light chain regulator STK17B (also known as DRAK2, implicated in several spinocerebellar ataxias [254, 255]), the cytoskeletal transport vesicle component TMED3 [256], the vesicle dynamics modulator SCRN1 [257], and the microtubule polymerization regulator DCLK1.

Importantly, downregulation of IGF1R as insulin-like growth factor 1 receptor tyrosine kinase was also documented. This observation is in line with the known Ataxin-2 modulation of MTOR growth signals, and it is noteworthy that a loss of function in IGF1R also leads to intrauterine lethality [258]. The reduction of the mitochondrial deacylase SIRT5 as metabolism modulator in dependence on nutrient availability [259–261] also suggests altered MTOR signals. MTOR can localize at lysosomes to sense nutrient availability and is coregulated with lysosomal enzymes, explaining the downregulation of MANBA as lysosomal mannosidase beta A, which is particularly abundant in adherent fibroblasts.

Finally, the proteome profile also exhibited downregulations of two factors implicated in inflammatory responses to bacterial infections or mitochondrial dysfunction. PYCARD contributes to innate immune response as integral adapter in inflammasome assembly which activates caspase-1 leading to secretion of pro-inflammatory cytokines [262–264]. NMES1 (aka COXFA4L3, MOCCI, or C15ORF58 in human) consists of only 83 amino acids that are localized to intracellular organelles that contain DNA and RNA in high concentration (nucleus and mitochondria). It is paralogous to mitochondrial respiratory chain factor NDUFA4 and was found to modulate cytochrome C oxidase during inflammation, so it is also known as Antiviral Mitochondrial Stress Response factor [265–268]. Via this modulation, it also induces stress-dependent lysosomal-autophagosomal degradation, upregulates glutathione as antioxidative protection factor, and enhances survival [269].

Thus, the proteome profile of MEF cells shows that the loss of ATXN2L leads to the depletion of its protein interactors NUFIP2 and SYNE2, and to strong indirect downstream protein regulations in the cytoskeletal transport machinery, in MTOR-dependent growth pathways, and in inflammation responses, but not in the RNA processing apparatus.

### Atxn2-CAG100-KnockIn pathogenesis involves protein accumulations

Current literature about the ATXN2 aggregation process that triggers neurodegeneration in SCA2 and ALS has paid much attention to the observation that TDP-43 is sequestrated as a factor that is essential for embryonic life [9, 12, 131, 270], and suggested the importance of abnormal ribonucleoprotein interactions and RNA toxicity in these diseases [271–273]. It was not appreciated that significant accumulation affects also the ribonucleoprotein ATXN2L as an abundant ATXN2 interactor, which is also essential for embryonic life [106], together with the ribonucleoprotein NUFIP2 and probably DDX6 as well. This sequestration is likely to trigger a partial loss-of-function for these ribonucleoproteins, thus impairing neuronal maintenance and survival. Moreover, the novel findings about accumulations of SYNE1 and DCLK1 protein in the aged spinal cord of *Atxn2*-CAG100-KIN mice provide additional evidence that altered cytoskeletal dynamics may contribute to the disease process in SCA2 and ALS. This concept is in agreement with recent discoveries of ALS genes that function in cytoskeletal integrity and axonal transport [274–276].

## Conclusions

The present study provides evidence that the poorly characterized ribonucleoprotein ATXN2L associates with numerous other ribonucleoproteins, of which only NUFIP2 depends on its presence. Downstream indirect consequences of ATXN2L loss have been shown to affect the cytoskeletal dynamics, mTOR-dependent growth pathways, and inflammation responses. The polyQ expansion of ATXN2 leads to an excess accumulation of ribonucleoproteins ATXN2L / NUFIP2 and of cytoskeletal factors SYNE1 / DCLK1, a finding with relevance to neurodegenerative diseases like SCA2 and ALS.

## Institutional review board statement

The study was conducted in accordance with the Declaration of Helsinki. The animal study protocol was approved by the Regierungspräsidium Darmstadt (V54-19c20/15-FK/1083, on 11 March 2019).

## CRediT authorship contribution statement

Conceptualization, S.G. and G.A.; methodology, G.K., D.M., L.-E.A.-M., A.R.K. and S.G.; validation, G.K., L.-E.A.-M. and A.R.K.; formal analysis, J.K., L.-E.A.-M. and G.A.; investigation, J.K., G.K., D.M., L.-E.A.-M. and A.R.K.; resources, J.K., N.-E.S. and S.G.; data curation, D.M. and G.A.; writing—original draft preparation, J.K., L.E.A.-M. and G.A.; writing—review and editing, J.K., L.-E.A.-M., N.-E.S., A.R.K., D.M. and G.A.; visualization, J.K., L.-E.A.-M., A.R.K. and G.A.; supervision, S.G. and G.A.; project administration, G.A.; funding acquisition, G.A. All authors have read and agreed to the published version of the manuscript.

## Acknowledgements

We thank Beata Lukaszewska-McGreal for proteome sample preparation and the Max Planck Society for support.

## Data availability

All proteome data were deposited in PRIDE with accession numbers PXD061048, and PXD061161.

## Supplementary material

**Figure S1.**
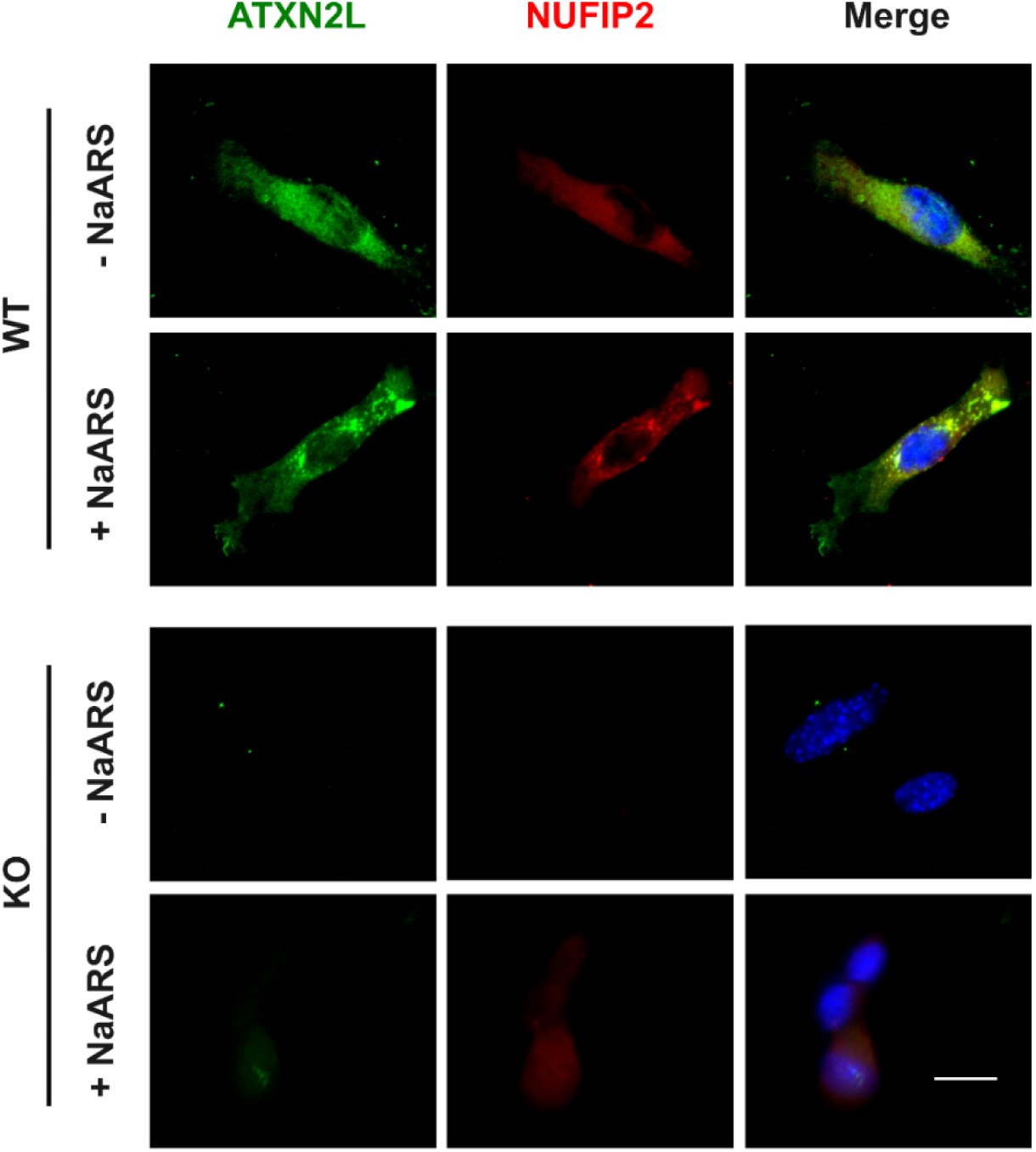
Triple immunofluorescence of ATXN2L (green color) and NUFIP2 (red color) in wildtype (WT) versus ATXN2L-null (KO) mouse embryonic fibroblasts show anti-ATXN2L positive foci colocalizing with NUFIP2 signals in stress granules induced by NaARS administration in mouse embryonic fibroblasts cultures. Scale bar represents 20 μm.

**Figure S2.**
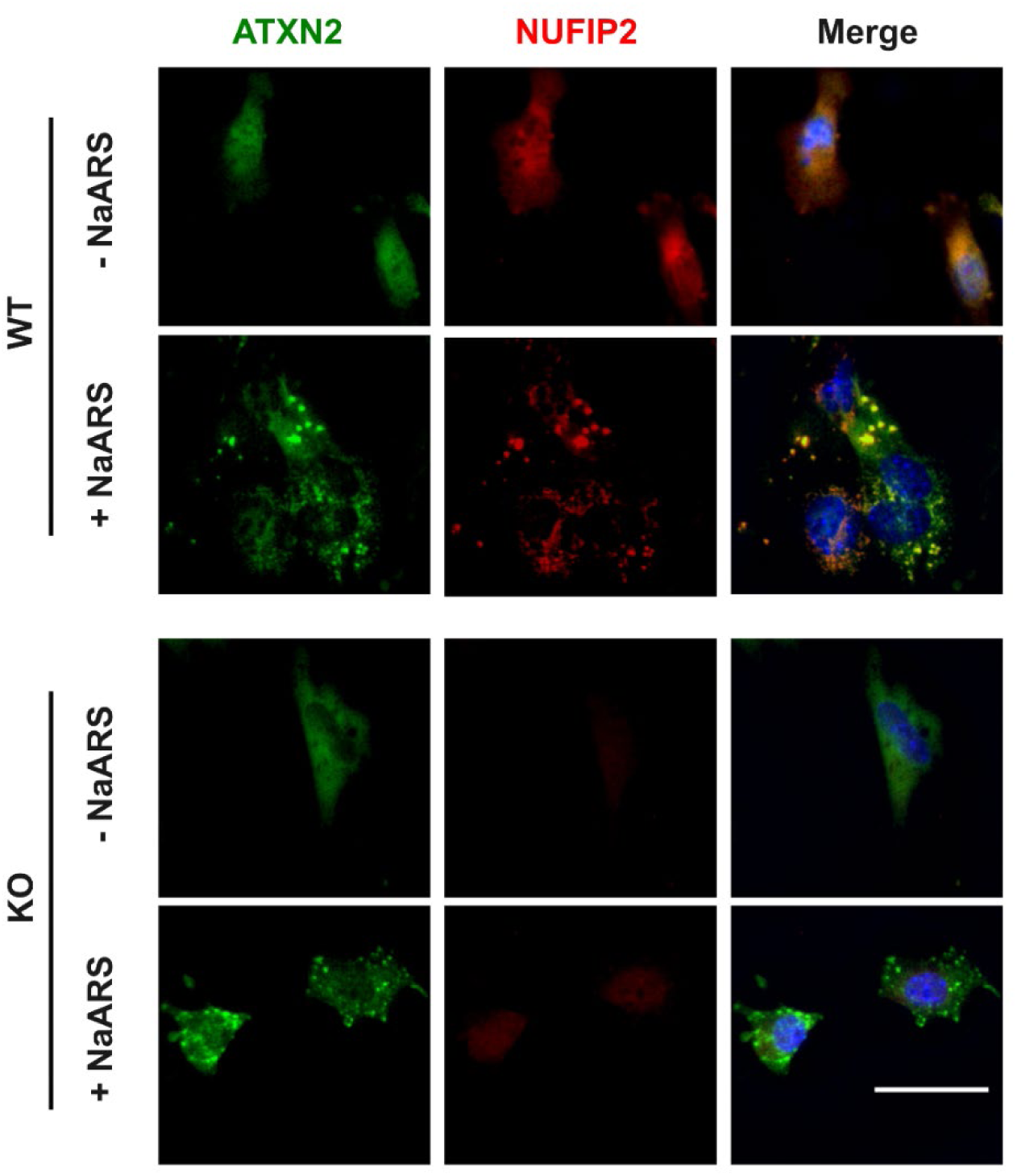
Triple immunofluorescence of ATXN2 (green color) and NUFIP2 (red color) in wildtype (WT) versus ATXN2L-null (KO) mouse embryonic fibroblasts show anti-ATXN2 positive foci colocalizing with NUFIP2 signals in stress granules induced by NaARS administration in mouse embryonic fibroblasts cultures. Scale bar represents 50 μm.

**Table S1: co-IP mass spectrometry quantifications.** Mass spectrometry identification and quantification files of proteome profiling of co-immunoprecipitates in MEF without stress or after NaARS stress. ATXN2L and its known interactor proteins are highlighted in yellow cells; protein abundance is illustrated by red background for high values and ratios above 1, while blue background illustrate ratios below 1; very low ratios are shown in dark blue.

**Table S2: Global proteome mass spectrometry quantifications.** Mass spectrometry identification and quantification files of proteome profiling of WT versus *Atxn2l-*null MEF. ATXN2L and its known interactor proteins are highlighted in yellow cells; protein abundance is illustrated by red background for high values and ratios above 1, while blue background illustrate ratios below 1; very low ratios are shown in dark blue.

